# Cortical interneuron loss and seizure generation as novel clinically relevant disease phenotypes in *Cln2^R207X^* mice

**DOI:** 10.1101/2022.03.11.483984

**Authors:** Keigo Takahashi, Elizabeth M. Eultgen, Sophie H. Wang, Nicholas R. Rensing, Hemanth R. Nelvagal, Joshua T. Dearborn, Mark S. Sands, Michael Wong, Jonathan D. Cooper

**Affiliations:** Department of Pediatrics, Washington University in St Louis, School of Medicine, St Louis, MO, USA; Department of Genetics, Washington University in St Louis, School of Medicine, St Louis, MO, USA; Department of Neurology, Washington University in St Louis, School of Medicine, St Louis, MO, USA; Department of Medicine, Washington University in St Louis, School of Medicine, St Louis, MO, USA

**Keywords:** Batten disease, CLN2 disease, neuronal ceroid lipofuscinosis, GABAergic interneurons, seizure generation, glial activation, neurodegeneration

## Abstract

**Aims:** CLN2 disease is a fatal inherited childhood neurodegenerative disorder. Although a disease-modifying therapy now exists, a fundamental lack of understanding of disease pathogenesis has hampered development of more effective therapies. To better understand the cellular pathophysiology of CLN2 disease, we investigated the nature and progression of neuropathological and neurological changes in the recently generated *Cln2^R207X^* mouse.

**Methods:** We have detailed microglial activation, astrogliosis, cytokine and chemokine expression, and neuron loss across the forebrain and spinal cords of *Cln2^R207X^* mice, along with quantitative gait analysis. We also performed long-term electroencephalography (EEG) recordings to characterize seizure activity, a clinically-relevant phenotype yet to be defined in any CLN2 disease model.

**Results:** Histology revealed early localized microglial activation months before neuron loss in the thalamocortical system and spinal cord, which was accompanied by astrogliosis. These pathological changes were more pronounced and occurred in the cortex before the thalamus or spinal cord. There were early-onset and progressive changes in the expression of specific chemokines and cytokines including IL-33, IP-10, and MIP-1α. Gait analysis revealed impaired performance only at disease end stage. EEG recordings revealed robust and progressive epileptiform activity from disease mid-stage including spontaneous seizures, which were accompanied by a profound loss of cortical GABAergic interneurons.

**Conclusions:** Our data reveal novel phenotypes in *Cln2^R207X^* mice that differ markedly in their timing and progression through the CNS from other NCL mouse models. Our findings provide new insights on CLN2 disease pathogenesis and clinically-relevant readouts for future therapeutic studies.

## Introduction

The neuronal ceroid lipofuscinoses (NCLs) are a group of neurodegenerative lysosomal storage disorders affecting children and young adults [1]. CLN2 disease, or classic late infantile neuronal ceroid lipofuscinosis, is one of the most common forms of NCL caused by a deficiency of tripeptidyl peptidase 1 (TPP1) [2–4]. Patients with CLN2 disease develop symptoms typically between 2 and 4 years of age including seizures and ataxia, often accompanied by speech delay [5, 6]. These are followed by progressive psychomotor deterioration and if untreated these children die during their teenage years [7]. A disease-modifying enzyme replacement therapy (ERT) for CLN2 disease is now available, and appears to slow disease progression [8, 9]. However, ERT requires invasive bi-weekly infusions and is incompletely effective. Therefore, there is still a need to develop less invasive and more effective therapies for CLN2 disease.

Progress towards this goal has been hampered by a fundamental lack of knowledge of how TPP1 deficiency affects the CNS. The histopathologic outcome measures used in NCL preclinical studies typically include storage material accumulation, glial activation and neuron loss [10, 11]. The onset and progression of these phenotypes has been characterized in detail in mouse models of multiple forms of NCL. These studies have revealed temporal and regional differences between the NCLs [12–19]. Such data have provided important information about the potential role for glia in NCL pathogenesis, and the anatomical regions that need to be targeted therapeutically. Although similar neuropathological phenotypes are evident in CLN2 knockout mice and have been used to judge efficacy in preclinical studies [20–22], the regional and cellular specificity of these phenotypes and how they progress over time has yet to be determined. Furthermore, although children with CLN2 disease usually first present with pronounced seizures, the seizure phenotype of CLN2 mice has yet to be explored.

We have addressed both of these knowledge gaps in a recently generated *Cln2^R207X^* mouse that models the most common human disease-causing mutation in the *CLN2* gene [23]. Our data reveal a range of pronounced differences in the staging of CLN2 neuropathology compared to other forms of NCL, and a robust spontaneous seizure phenotype associated with cortical interneuron loss. These data provide key information for the effective targeting of pre-clinical interventions, as well as a robust clinically-relevant outcome measure for judging their efficacy.

## Materials and Methods

### Mice

*Cln2^R207X^* mice were first generated by Geraets et al. [23]. *Cln2^R207X^* and wild-type (WT) mice were maintained separately on a congenic C57Bl/6J background and housed in an animal facility at Washington University School of Medicine (St. Louis, MO) under a 12hr light/dark cycle, and provided food and water *ad libitum*. All procedures were performed in accordance with NIH guidelines under a protocol approved by the Institutional Animal Care and Use Committee (IACUC) at Washington University School of Medicine. Unless otherwise stated, n=6 (3 male, 3 female) mice of each genotype were used, except for gait analysis/EEG recording (n=10; 5 male, 5 female) and RT-qPCR/neurotransmitter analysis (n=4; 2 male, 2 female).

### Tissue processing and Nissl staining

Processing of both forebrain and lumbar spinal cord tissues and Nissl staining were performed, as described previously [18,24,25].

### Immunohistochemistry and imaging

A one-in-six series of coronal forebrain sections and a one-in-forty-eight series of 40µm coronal spinal cord sections from each mouse were stained on slides using a standard immunofluorescence protocol [19, 26] for the following antibodies: rabbit anti-GFAP 1:1000, DAKO, rat anti-CD68 1:400, Bio-Rad, rabbit anti-SCMAS 1:400, abcam, rat anti-MHC-II 1:200, Invitrogen, rat anti-Clec7a 1:200, InvivoGen, rabbit anti-parvalbumin (PV) 1:200, Swant, mouse anti-GAD67 1:200 EMD Millipore. To visualize GABAergic interneurons, a one-in-six series of coronal sections were stained on slides using a modified peroxidase protocol [18, 24]. After quenching endogenous peroxidase activity in 1% H_2_O_2_ and blocking in 15% normal swine serum (Vector) for 1 hour, sections were incubated in primary antibodies for interneuron markers (rabbit anti-PV, 1:1000, rabbit anti-calbindin (CB), 1:1000, and rabbit anti-calretinin (CR), 1:1000, all from Swant) overnight at 4°C. Subsequently, sections were incubated for 2 hours in biotinylated swine anti-rabbit secondary antibody (1:200, Dako) followed by 2-hour incubation in Vectastain ABC (avidin-biotin, 1:200, Vector), and immunoreactivity visualized using 0.05% DAB (Sigma) and 0.005% H_2_O_2_. All images were taken on a Zeiss *AxioImager Z1* microscope using *StereoInvestigator* (MBF Bioscience) software or a Zeiss LSM880 Confocal Laser Scanning Microscope with *Airyscan* and ZEN 2 (blue edition, Zeiss) software.

### Quantitative thresholding image analysis

To quantify AFSM accumulation and glial activation (GFAP-positive astrocytes and CD68-, MHC-II-, or Clec7a-positive microglia), semiautomated thresholding image analysis was performed as described previously [19, 26]. This involved collecting slide-scanned images at 10x magnification (Zeiss Axio Scan Z1 Fluorescence Slide Scanner) from each animal. Images were subsequently analyzed using *Image-Pro Premier* (Media Cybernetics) using an appropriate threshold that selected the foreground immunoreactivity above background.

### Neuron counts

Unbiased design-based optical fractionator counts of Nissl-stained neurons and GABAergic interneurons were performed using *StereoInvestigator* (MBF Bioscience) in a one-in-six series of forebrain hemisections and a one in forty-eight spinal cord hemisections, as described previously [18,19,24]. The following sampling scheme and objective lens were applied to each region of interest. Specific sizes of sampling grid and counting frame and appropriate objectives were used for each region and staining, as detailed in **Supporting Methods**.

### Total RNA extraction, cDNA synthesis, and RT-qPCR

Total RNA was extracted from forebrain homogenates from each animal and purified using TRIzol (Thermo Fisher), as previously described [27]. Subsequently, cDNA was synthesized using Random Primers (Invitrogen) and SuperScript II Reverse Transcriptase (Invitrogen) according to the manufacturer’s protocol. All RT-qPCR reactions for each gene of interest were performed in triplicate. RT-qPCR was performed using SYBR Green PCR Master Mix (Applied Biosystem) with primers whose sequences are listed in **Supporting Methods**. 2^−ΔΔCt^ of gene expression was analyzed for each gene and normalized with the reference gene, glyceraldehyde 3-phosphate dehydrogenase (GAPDH).

### Cytokine and chemokine profile analysis

Cytokine and chemokine profiling in proteins extracted from forebrain homogenates was performed using an Affymetrix multiplex assay, as described previously [19]. Customized Procartaplex Luminex Assay 24 plex for CRP, ENA-78 (CXCL5), Eotaxin (CCL11), GRO α (KC/CXCL1), IFN-α, IFN-β, IFN-γ, IL-1α, IL-1β, IL-2, IL-4, IL-6, IL-10, IL-33, IP-10 (CXCL10), MCP-1 (CCL2), MCP-3 (CCL7), M-CSF, MIP-1α (CCL3), MIP-1β (CCL4), MIP-2α (CXCL2), RANTES (CCL5), TNF-α, and VEGF-A were run in duplicate.

### Tissue cultures and pharmacological stimulation

Both primary astrocytes and microglia were generated from mixed glial cells isolated from postnatal day 1-5 (P1-5) WT or *Cln2^R207X^* mouse cortices as previously described [28, 29]. Briefly, astrocytes were generated by shaking confluent mixed-glial cultures at 10-14 days *in vitro* (DIV) at 180 rpm overnight to remove microglia followed by 7 days of treatment with cytosine arabinoside (Ara-C, Sigma). To generate microglia, confluent mixed glial cultures at 14-18 DIV were subsequently incubated in Trypsin-EDTA (0.5mg/ml, Sigma) for 45-60 minutes until the astrocyte monolayer was removed. For pharmacological stimulation, astrocytes were stimulated with lipopolysaccharide (LPS, 1μg/ml, Sigma) and IFN-γ (800U/ml, PeproTech) for 48 hours, while microglia were stimulated with LPS alone for 6 hours [28, 29]. Both astrocytes and microglia for immunocytochemistry were plated onto poly-D-lysine (PDL) coated glass coverslips (Neuvitro Corporation) in 24 well plates. Cell culture experiments were repeated three times independently (n=3 per group).

### Immunocytochemistry and image analysis

Cultures were fixed in 4% PFA and stained using a standard immunocytochemistry protocol [28, 29] for astrocytes (rabbit anti-GFAP 1:500, Dako, mouse anti-α-tubulin 1:500, Sigma) and microglia (rabbit anti-Iba1, 1:500, FUJIFILM Wako). Images of 10 random fields per coverslip were taken at 20x magnifications. Among astrocytes visualized by α-tubulin staining, the number of GFAP-positive and negative astrocytes were manually counted. The cell surface of Iba1-positive microglia was outlined manually, and the aspect ratio and roundness were automatically calculated by *StereoInvestigator* (MBF Bioscience). All analyses were performed with three coverslips for each repeated experiment and reported as an average.

### Quantitative gait analysis

The CatWalk XT gait analysis system (Noldus Information Technology, Wagenigen, Netherlands) was used to study gait performance, as described previously [19, 30].

### Electroencephalography monitoring

WT and *Cln2^R207X^* mice underwent continuous video-EEG monitoring starting at 10 weeks of age, using standard methods for implanting epidural electrodes and EEG recording under isoflurane anesthesia, as previously described [31–33]. Burr holes for the frontal reference electrodes were made (anterior +0.5mm, lateral +/-0.5mm; bregma) and secured via screws. Two bilateral “active” recording electrodes were placed over the parietal cortex (posterior -2.5mm, lateral +/- 1.5; bregma) and a ground screw secured over the cerebellum (posterior -6.2mm, lateral +/- 0.5; bregma). At least 72 hours after recovering from surgery groups of four mice were placed in individual cages and connected via a custom flexible cable attached to the exposed pin header for recording. Continuous bilateral cortical video-EEG signals starting at 10 weeks of age were acquired using a referential montage using Stellate or LabChart (AdInstruments) acquisition software and amplifiers until death for *Cln2^R207X^* mice or the end of 20 weeks for WT mice. Signals were amplified at 10,000X with high-pass (0.5Hz) and low-pass (100Hz) filters applied. EEG signals were digitized at 250Hz and time-locked video EEG collected continuously.

### Neurotransmitter analysis

The concentration of cortical neurotransmitters including glutamate (Glu), glutamine (Gln), and gamma-aminobutyric acid (GABA) was measured by liquid chromatography with tandem mass spectrometry (LC-MS/MS). Mouse cortex samples were homogenized in water (10 mL/g tissue). All amino acids listed above were extracted from 20μL of homogenate with 200μL of methanol after addition of 20μL of internal standard solution containing Glu-d_3_ (500μg/mL), Gln-^13^C_5_ (300μg/mL), and GABA-d_6_ (300μg/mL). A seven-point standard curve was prepared in duplicate. The analysis of amino acids was performed on a Shimadzu 20AD HPLC system and a SIL-20AC autosampler coupled to 4000Qtrap mass spectrometer (AB Sciex) operated in positive multiple reaction monitoring (MRM) mode. Data processing was conducted with Analyst 1.6.3 (Applied Biosystems). Calibrators that deviate by more than 15% of nominal concentrations were excluded from construction of calibration curve, except that deviation of 20% was acceptable for the lower limit of quantification (LLOQ), which is the low end of calibration curve.

### Statistical analysis

All statistical analyses were performed using GraphPad Prism version 9.1.0 for MacOS (GraphPad Software, San Diego, CA). A multiple t-test with a post-hoc Holm-Šídák correction was used for histological data at multiple time points and cytokine and chemokine profiling data. Unpaired t-test was used for Nissl-stained neuron and interneuron counting within the hippocampus. A one-way ANOVA with a post-hoc Bonferroni correction was used for morphological analysis of tissue cultures and neurotransmitter analysis, and a two-way ANOVA with a post-hoc Bonferroni correction for gait analysis. A p-value of ≤0.05 was considered significant.

## Results

### Localized early microglial activation precedes astrogliosis in the *Cln2^R207X^* mouse

Both microglial activation and astrogliosis have previously been documented only at the disease end stage in *Cln2^R207X^* mouse brains [23]. To define the staging of these neuroimmune responses and their spatiotemporal relationship with neuron loss in *Cln2^R207X^* mice, we surveyed brain and spinal sections immunostained for multiple markers of microglial activation and astrogliosis. Thresholding image analysis revealed that a localized CD68-positive microglial activation starts within the primary motor (M1), somatosensory (S1BF), and visual (V1) cortices as well as the spinal ventral horn at 1 month. This increase in CD68 immunoreactivity appeared to be most pronounced within the somatosensory thalamocortical system including S1BF and ventral posterior thalamic nuclei (VPL/VPM) compared with other regions (Figure 1A-B). Different states of microglial activation resembling the M1/M2 polarization of macrophages, and the expression of disease-associated microglia (DAM) have both been recently implicated in several neurodegenerative diseases [34–36]. Therefore, we additionally stained *Cln2^R207X^* forebrain sections for major histocompatibility class II (MHCII, M1 marker) and C-type lectin domain containing 7a (Clec7a, M2 and DAM marker). MHCII-positive and Clec7a-positive microglia were observed in the VPL/VPM of *Cln2^R207X^* mice only at the disease end stage (Figure S1A), suggesting the presence of different states of microglial activation between early and late stage of CLN2 disease.

**Figure 1.**
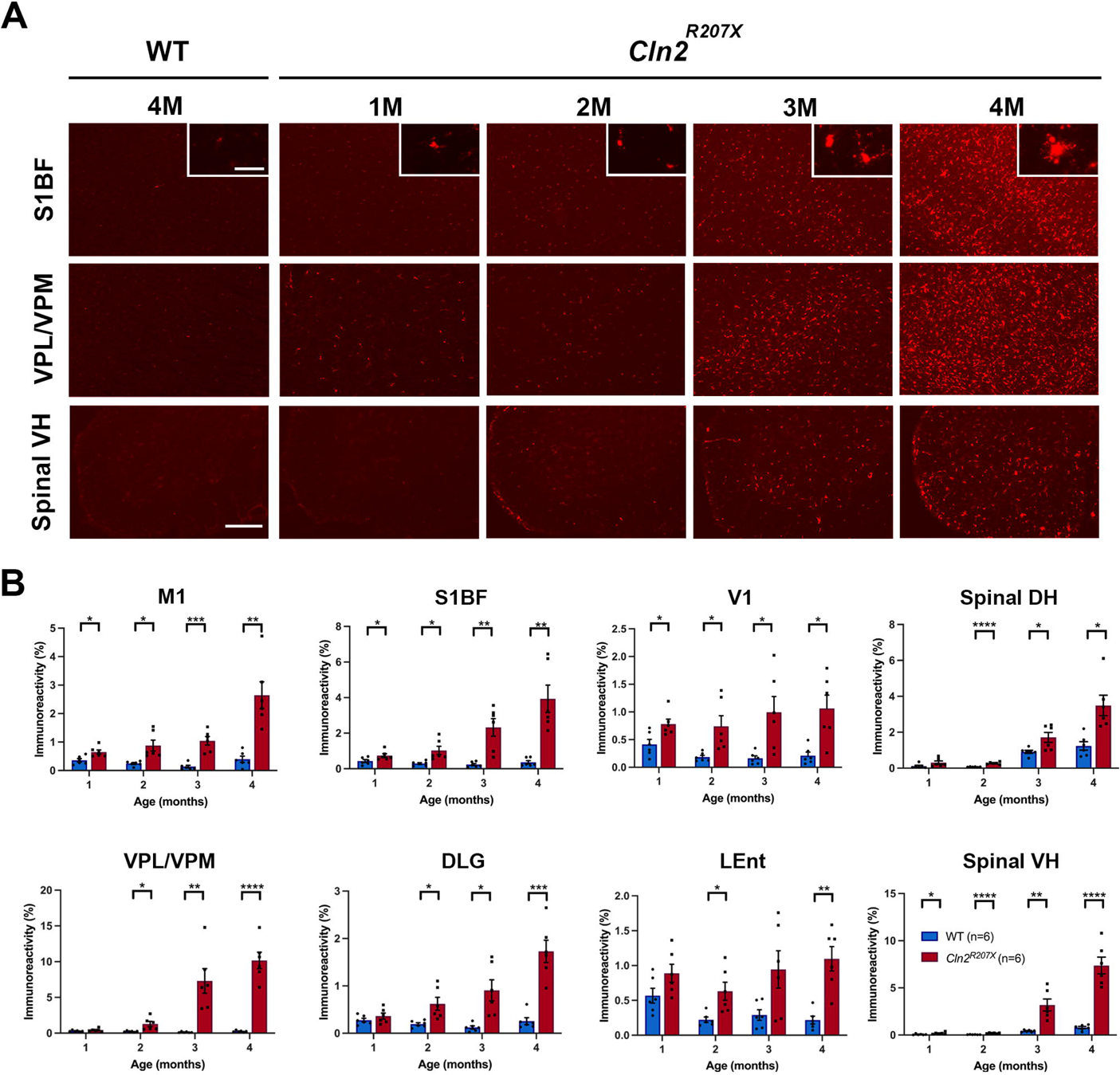
*Cln2^R207X^* mice show localized early microglial activation. (*A*) Immunostaining for the microglial marker CD68 (red) reveals the pronounced increase in abundance and staining intensity of CD68 immunoreactivity with disease progression in the primary somatosensory cortex (S1BF), ventral posterior nuclei of thalamus (VPL/VPM), and lumbar spinal ventral horn (VH) compared with WT control mice. Corresponding changes in microglial morphology are revealed in higher power inserts [Scale bars = 200 μm and 20 μm (*Insets*)]. (*B*) Quantitative thresholding image analysis of CD68 immunoreactivity confirms the progressive microglial activation in *Cln2^R207X^* mice (red bars) beginning in the primary motor cortex (M1), S1BF, primary visual cortex (V1), and spinal VH at 1 months of age, followed by microglial activation in the VPL/VPM, dorsolateral geniculate nucleus (DLG), lateral entorhinal cortex (LEnt), and lumbar spinal dorsal horn (DH) starting at 2 months of age compared with age-matched WT control mice (blue bars). Dots represent values from individual animals. \**P* < 0.05, \*\**P* < 0.01, \*\*\**P* < 0.001, \*\*\*\**P* < 0.0001, multiple t-test with Holm-Šídák correction. Values are shown as mean ± SEM (n = 6 mice per group).

Staining for glial fibrillary acidic protein (GFAP, astrogliosis marker) revealed an increase in GFAP immunoreactivity that became significant as early as 2 months in *Cln2^R207X^* brains and 3 months in spinal cords (Figure 2A-B), indicating an onset of astrogliosis that was later than that of microglial activation. Similar to CD68, GFAP immunoreactivity was higher within S1BF and VPL/VPM than in other regions (Figure 2A-B). Recently, the concept of A1/A2 astrocytes, in which A1 is neurotoxic and A2 is neuroprotective has been proposed [37]. Therefore, we investigated expression levels of 14 different genes known to be specific to A1 or A2 astrocytes in *Cln2^R207X^* forebrains by RT-qPCR. While Serping1 (A1-specific) and Clcf1 (A2-specific) were significantly elevated at 3 months, those genes were rather suppressed at 4 months. In contrast, C3 and H2-D1 (A1-specific) and Cd14 (A2-specific) were significantly elevated (Figure S1B), demonstrating an expression profile of A1/A2-specific genes in *Cln2^R207X^* forebrains that differs from other neurodegenerative diseases.

**Figure 2.**
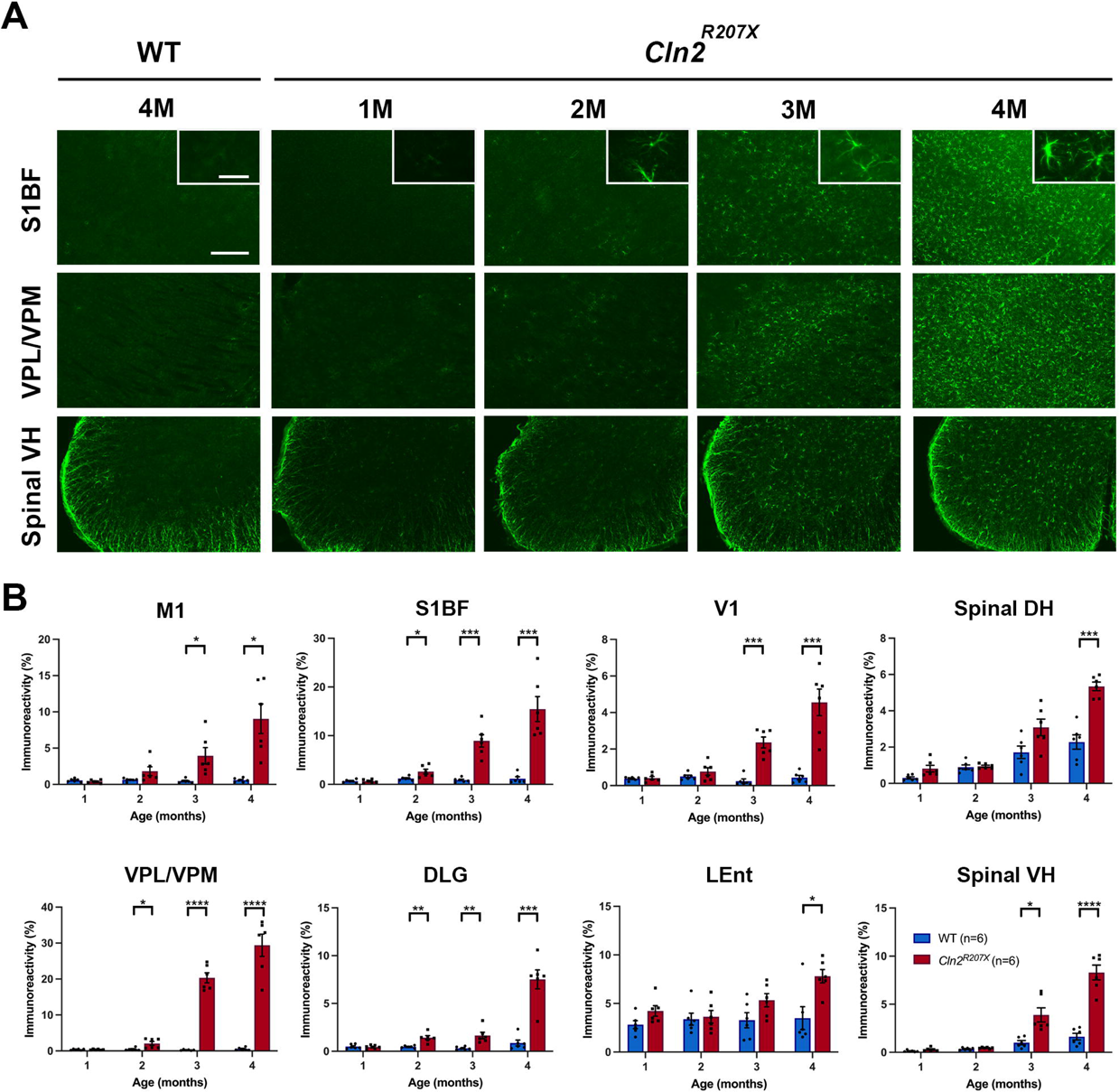
*Cln2^R207X^* mice show relatively delayed onset of astrogliosis. (*A*) Immunostaining for the astrocyte marker glial fibrillary associated protein (GFAP, green) reveals the marked increase in the abundance of GFAP positive cells and the intensity of GFAP immunoreactivity with disease progression in the primary somatosensory cortex (S1BF), ventral posterior nuclei of thalamus (VPL/VPM), and lumbar spinal ventral horn (VH) compared with WT control mice. Corresponding changes in astrocyte morphology are revealed in higher power inserts [Scale bars = 200 μm and 20 μm (*Insets*)]. (*B*) Quantitative thresholding image analysis of GFAP immunoreactivity confirms the progressive nature of astrogliosis at 12-, 24-, 36-and 48-week-old time points in in the primary motor cortex (M1), S1BF, primary visual cortex (V1), VPL/VPM, dorsolateral geniculate nucleus (DLG), lateral entorhinal cortex (LEnt), lumbar spinal dorsal horn (DH), and lumbar spinal VH as early as 2 months of age in *Cln2^R207X^* mice (red bars) compared with age-matched WT control mice (blue bars). Dots represent values from individual animals. \**P* < 0.05, \*\**P* < 0.01, \*\*\**P* < 0.001, \*\*\*\**P* < 0.0001, multiple t-test with Holm-Šídák correction. Values are shown as mean ± SEM (n = 6 mice per group).

### Distinct neuroinflammatory environment in *Cln2^R207X^* mouse forebrains

Having documented the staging of microglial activation and astrogliosis, we further investigated the nature of the neuroimmune response in *Cln2^R207X^* forebrains by measuring 23 chemokines and cytokines. There was a significant increase in IL-33 starting at 1 month and significant increases in IP-10 (CXCL10) and MIP-1α (CCL3) starting at 2 months compared with wild-type controls (Figure 3). Increases in MCP-1 (CCL2), MCP-3 (CCL7), M-CSF (CSF1), VEGF-A, IL-4, MIP-2 (CXCL2) were observed at 3 months. Finally, a significant increase in RANTES (CCL5), and decrease in ENA-78 (CXCL5) were observed at 4 months (Figure 3). There were no significant changes in a number of pro- and anti-inflammatory molecules including IL-1α/β, TNF-α, IFN-γ, IFN-α, IFN-β, IL-2, IL-6, IL-10, CXCL1, and CRP at any time point (Figure S2).

**Figure 3.**
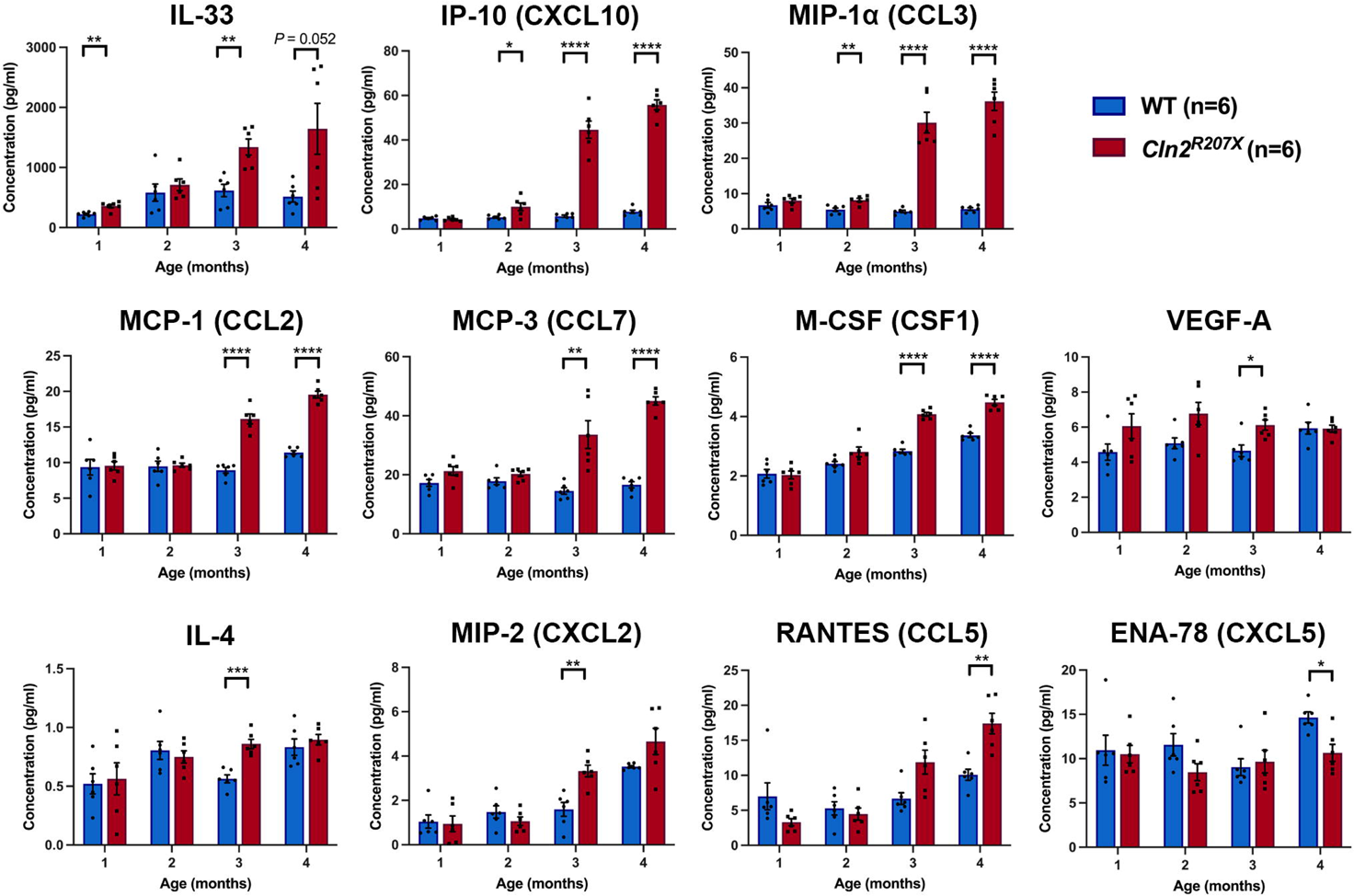
*Cln2^R207X^* mouse forebrains show distinct neuroinflammatory changes. Observed concentrations (pg/ml) of multiple cytokines and chemokines measured in forebrain homogenates reveal significant elevation of IL-33 at 1 month, IP-10 (CXCL10) and MIP-1a (CCL3) at 2 months, MCP-1 (CCL2), MCP-3 (CCL7), M-CSF (CSF1), VEGF-A, IL-4, and MIP-2 (CXCL2) at 3 months, and VEGF-A (CCL5) at 4 months of age and significant reduction of ENA-78 (CXCL5) at 4 months of age in *Cln2^R207X^* forebrains (red bars) compared with age-matched WT controls (blue bars). Dots represent values from individual animals. \**P* < 0.05, \*\**P* < 0.01, \*\*\**P* < 0.001, \*\*\*\**P* < 0.0001, multiple t-test with Holm-Šídák correction. Values are shown as mean ± SEM (n = 6 mice per group).

### Normal microglial responses, but compromised astrocytic responses to pharmacological stimulation *in vitro*

To further investigate the influence of the *Cln2^R207X^* mutation upon glial function, we compared the reactivity and response of microglia and astrocytes from *Cln2^R207X^* and WT mice to various stimuli *in vitro*. We first assessed the ability of Iba1-positive primary microglia to undergo a morphological transformation in response to pharmacological stimulation with LPS. Morphological analysis of *Cln2^R207X^* microglia under basal conditions revealed no change in aspect ratio and roundness compared with WT microglia, indicative of a normal basal activated state (Figure S3A-B). Also, following LPS treatment, both WT and *Cln2^R207X^* microglia responded similarly with a significantly higher aspect ratio and roundness compared with their basal morphologies (Figure S3A-B).

Next, we assessed GFAP immunoreactivity in primary astrocytes under basal conditions and following pharmacological stimulation with LPS and IFN-γ. Under basal conditions, there was no significant difference in the proportion of GFAP-positive astrocytes between WT and *Cln2^R207X^* (Figure S3C-D). Following LPS/IFN-γ treatment, while the fraction of GFAP-positive cells in WT astrocytes was significantly increased, those of *Cln2^R207X^* showed no change (Figure S3C-D).

### Neuron loss occurs in the forebrain of *Cln2^R207X^* mice before the spinal cord

To define the onset and progression of neuron loss and its spatiotemporal relationship with microglial activation and astrogliosis, we performed unbiased stereological counting of Nissl-stained neurons in the thalamocortical system and spinal cords of *Cln2^R207X^* mice. A significant reduction in the number of Nissl-stained neurons within S1BF lamina V was first present at 3 months and in VPL/VPM at 4 months (Figure 4). In contrast, significant loss of spinal cord dorsal horn neurons in lamina III-V was only present at 4 months, with no significant loss of neurons in the dorsal horn at any age (Figure 4). No significant changes were observed in CA1 hippocampal neurons (Figure S4), indicating the distinct anatomical distribution of Nissl-stained neuron loss in *Cln2^R207X^* mice.

**Figure 4.**
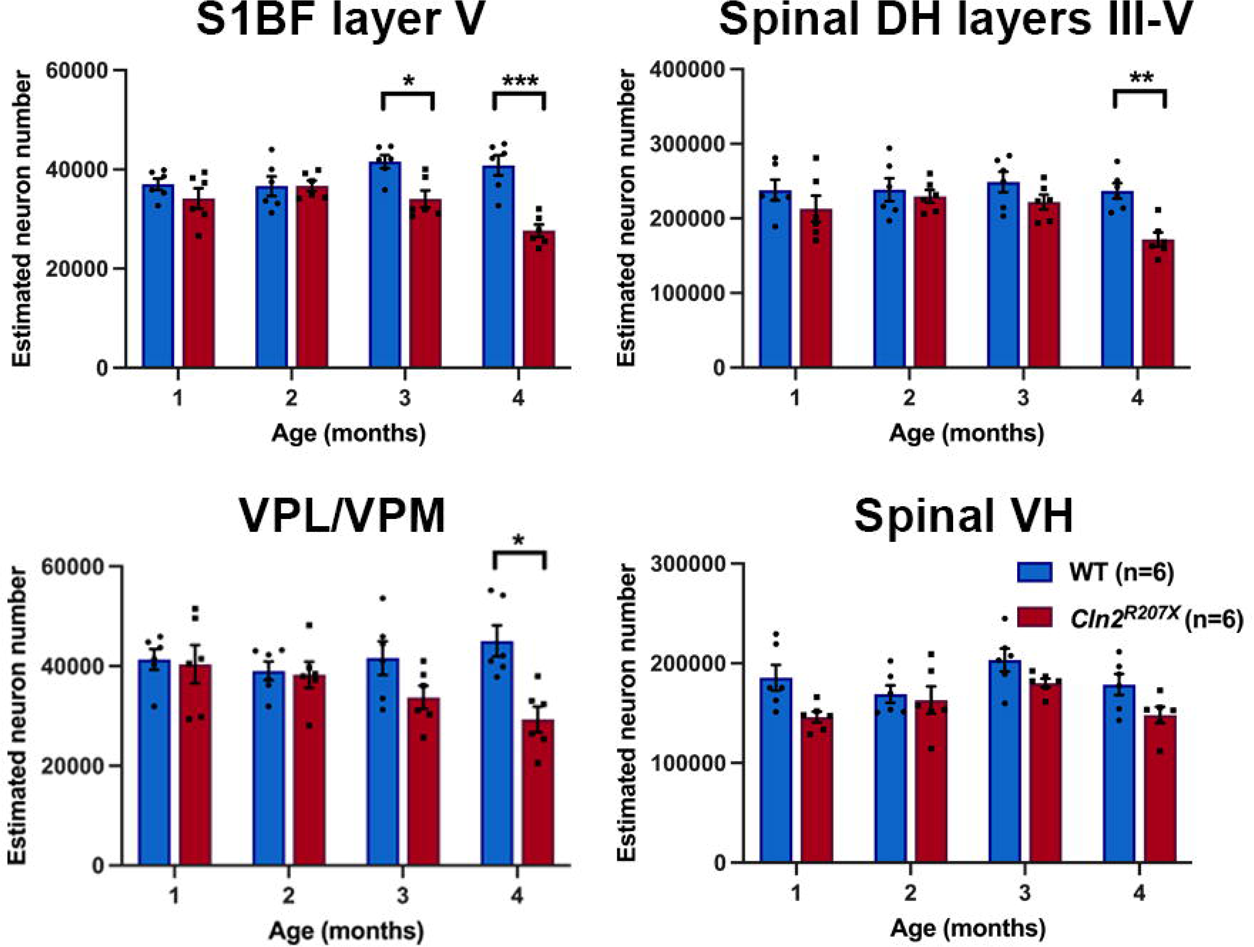
*Cln2^R207X^* mice show distinct staging of Nissl-stained neuron loss in the CNS. Unbiased stereological counting reveals loss of Nissl-stained neurons becomes significant within primary somatosensory cortex (S1BF) lamina V at 3 months and within ventral posterior nuclei of thalamus (VPL/VPM) and lumbar spinal dorsal horn (DH) lamina III-V at 4 months of age in *Cln2^R207X^* mice (red bars) compared with age-matched WT control mice (blue bars), whereas there is no significant difference between genotypes in lumbar spinal cord ventral horn (VH). Dots represent values from individual animals. \**P* < 0.05, \*\**P* < 0.01, \*\*\**P* < 0.001, \*\*\*\**P* < 0.0001, multiple t-test with Holm-Šídák correction. Values are shown as mean ± SEM (n = 6 mice per group).

### Early storage material accumulation is widespread storage material in the CNS of *Cln2^R207X^* mice

Storage material accumulation within neurons and glia is a characteristic CNS neuropathological hallmark of the NCLs [10, 11]. Although it does not appear to be toxic, storage burden is a useful quantitative outcome measure for assessing therapeutic efficacy. Staining for subunit-c of mitochondrial ATPase (SCMAS), a major storage component in CLN2 disease, revealed a significant increase in SCMAS immunoreactivity as early as 1 month across all analyzed regions of the *Cln2^R207X^* CNS (Figure 5A and B). Given such an early onset and a pronounced degree of SCMAS accumulation, we also analyzed neonatal *Cln2^R207X^* brains. Thresholding analysis confirmed significant accumulation of SCMAS within most brain regions at P1 (Figure 5B and S5).

**Figure 5.**
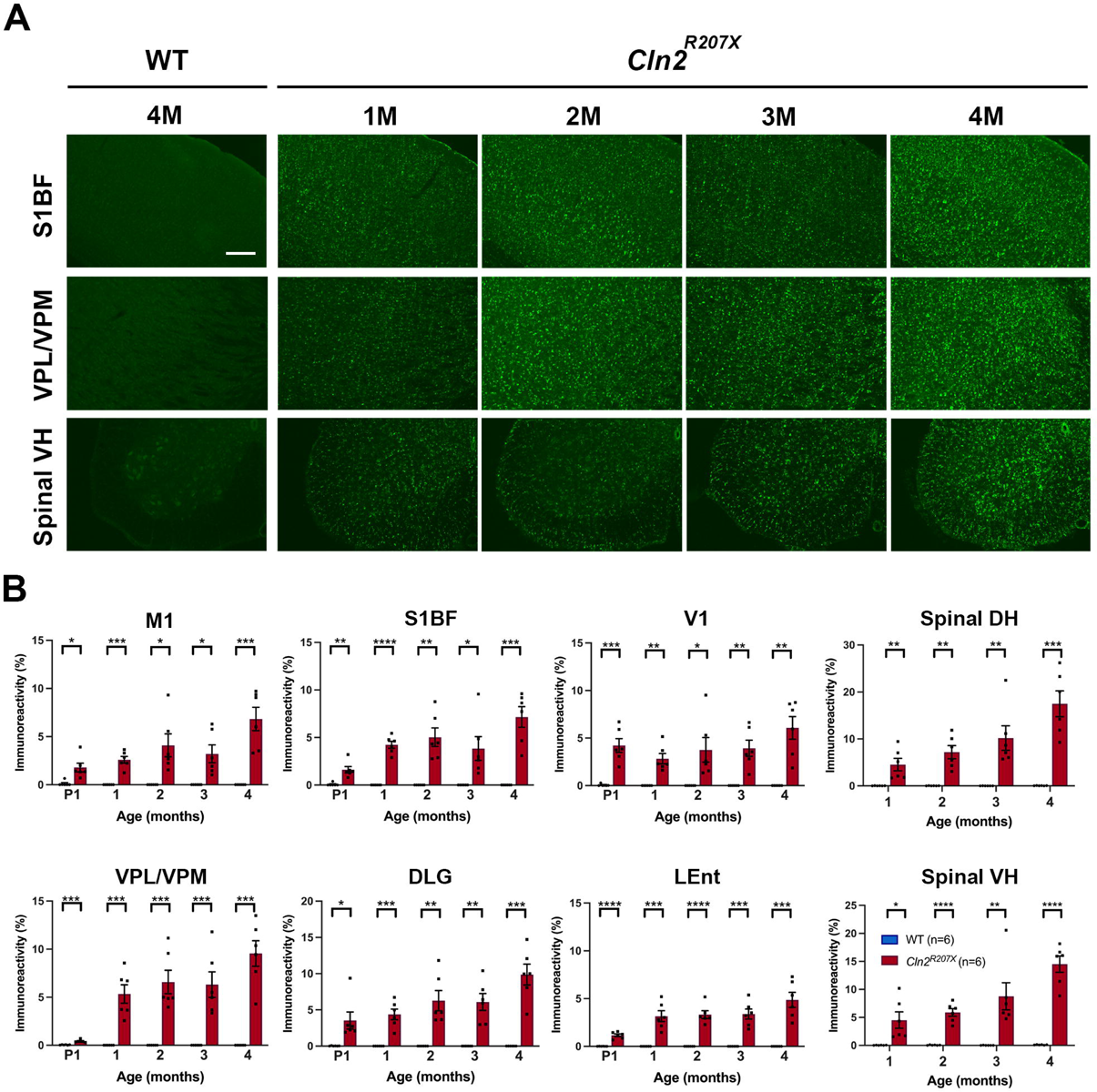
*Cln2^R207X^* mice show early widespread lysosomal storage material accumulation across the CNS. (*A*) Immunostaining for subunit c of mitochondrial ATP synthase (SCMAS, green) reveal the widespread increase in SCMAS accumulation in the primary somatosensory cortex (S1BF), ventral posterior nuclei of thalamus (VPL/VPM), and lumbar spinal ventral horn (VH) compared with WT control mice as early as 1 months. Scale bar = 200 μm. (*B*) Quantitative analysis of SCMAS immunoreactivity via thresholding image analysis in the same brain regions confirms the early onset of SCMAS accumulation across many CNS regions in *Cln2^R207X^* mice (red bars) compared with age-matched WT control mice (blue bars). Dots represent values from individual animals. \**P* < 0.05, \*\**P* < 0.01, \*\*\**P* < 0.001, \*\*\*\**P* < 0.0001, multiple t- test with Holm-Šídák correction. Values are shown as mean ± SEM (n = 6 mice per group).

### Late-onset gait abnormalities in *Cln2^R207X^* mice

Gait disturbances are one of major symptoms of CLN2 disease and clinically used to evaluate disease progression in patients [38, 39]. Gait abnormalities in *Cln2^-/-^* mice at disease end stage have been previously documented using a simple but time-consuming and rudimentary footprint analysis [21]. Here, we comprehensively quantified the nature and progression of gait abnormalities in *Cln2^R207X^* mice using a CatWalk^XT^ apparatus, a fully automated and objective gait analysis system. *Cln2^R207X^* mice exhibited significantly shorter average stride length and average swing duration at 4 months and spent more time supporting themselves upon three and four feet from 3-4 months (Figure 6), suggesting late-onset of locomotor deficits. Nevertheless, many other gait parameters such as average body speed, cadence, average stand, and average step cycle were not significantly altered in *Cln2^R207X^* mice (Figure 6), suggesting relatively mild and late-onset gait abnormalities in *Cln2^R207X^* mice.

**Figure 6.**
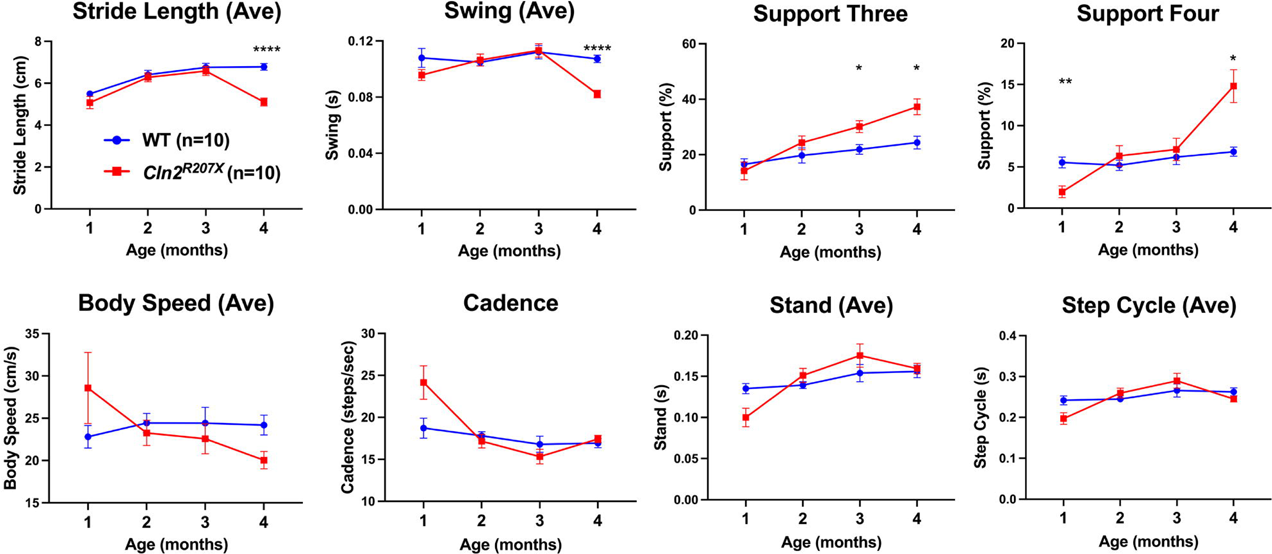
*Cln2^R207X^* mice show late-onset gait abnormalities. CatWalk^XT^ gait analysis reveals significantly shorter average stride length and average swing duration at 4 months and higher proportion of steps supported by three or four feet at 3-4 months of age in *Cln2^R207X^* mice (red lines) compared with age-matched WT control mice (blue lines). Other parameters including average body speed, cadence, average stand, and average step cycle showed no significant change between genotypes. \**P* < 0.05, \*\**P* < 0.01, \*\*\**P* < 0.001, \*\*\*\**P* < 0.0001, two-way ANOVA with Bonferroni correction. Values are shown as mean ± SEM (n = 10 mice per group).

### EEG monitoring reveals a pronounced seizure phenotype in *Cln2^R207X^* mice

We have noticed that *Cln2^R207X^* mice often die suddenly at around 4 months of age, especially during bedding changes or when transported between rooms. Considering that epileptic seizure is a major initial symptom, and is therapy-resistant in patients with CLN2 disease [5–7], we performed long-term video-EEG recordings to investigate if *Cln2^R207X^* mice also develop any seizure-related activity. WT control mice showed a normal EEG background activity consisting of mixed background EEG frequencies and no epileptiform activity (data not shown). At 10 weeks *Cln2^R207X^* mice exhibited a similar EEG background compared to WT mice, but at around 12 weeks of age they started to display epileptiform spikes and spike bursts superimposed on a mostly normal background (Figure 7A). A progressive increase in background spiking and clustering of spike bursts occurred over the 12-14 week period, and resulted in generalized EEG slowing, frequent spiking, bursting, and often burst-suppression activity by 16 weeks of age (Figure 7A). Starting at 14 weeks of age, *Cln2^R207X^* mice developed stereotypical spontaneous electrographic seizures involving a sudden tonic spike discharge which evolved in frequency and amplitude, and lasted an average of 34.5 seconds (Figure 7A-B). Clinically, seizures were typically characterized by head bobbing, rearing with forelimb clonus, and occasional generalized convulsive activity. These spontaneous seizures were present in 90% of *Cln2^R207X^* mice, with a median age at onset of 15 weeks (Figure 7D). Most *Cln2^R207X^* mice had repetitive seizures over a couple weeks, but some had a single seizure immediately prior to death. Notably, 70% of *Cln2^R207X^* mice died within 3 minutes of their last seizure (Figure 7C).

**Figure 7.**
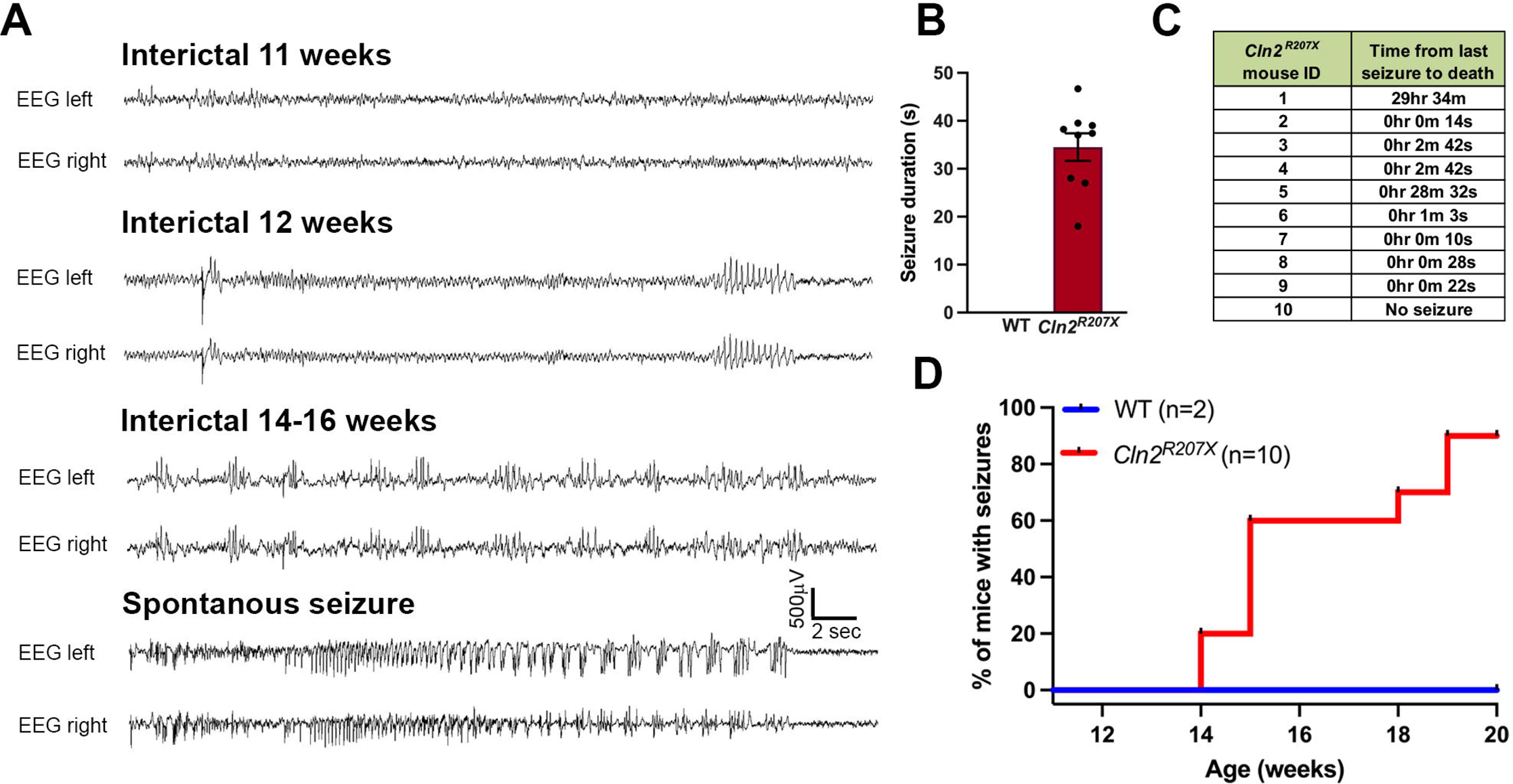
*Cln2^R207X^* mice show a pronounced seizure phenotype. Electroencephalography (EEG) recording reveals epileptiform interictal abnormality and spontaneous seizures in *Cln2^R207X^* mice. (*A*) Representative EEG traces at different time points in *Cln2^R207X^* mice. (*B*) Average duration of spontaneous seizures in *Cln2^R207X^* mice. (*C*) Interval between last seizure and death in *Cln2^R207X^* mice. (*D*) Percentage of mice displaying spontaneous seizure up to 20 weeks of age in WT (n=2) and *Cln2^R207X^* mice (n=10), with a median age at onset of 15 weeks.

### Loss of GABAergic interneurons precedes a generalized loss of neurons

GABAergic interneuron dysfunction and loss have been linked to epileptogenesis [40, 41]. To determine the relationship between altered EEG activity and potential interneuron loss we stained *Cln2^R207X^* forebrain sections for several interneuron-specific calcium-binding proteins including parvalbumin (PV), calbindin (CB), and calretinin (CR). Histological analysis revealed a progressive decline in the number of PV- and CB-positive interneurons as early as 2 months in the S1BF and from 3 months in the thalamic reticular nucleus (Figure 8A). The distribution of PV-positive fibers and terminals within S1BF was drastically diminished within lamina II-III of S1BF as early as 3 months (Figure 8C). This laminar specific effect within S1BF was precisely mirrored in the same sections counter-stained for glutamate decarboxylase 67 (GAD67), an enzyme that metabolizes glutamate into GABA (Figure 8C). The number of CR-positive interneurons was also significantly reduced at 1 and 4 months (Figure 8A), suggesting both developmental and degenerative changes in CR-positive interneurons. Consistent with counts of Nissl stained CA1 neurons, no changes were observed in the number of any interneuron population within the hippocampus (Figure S4). To further investigate the nature of GABAergic dysfunction, we quantified concentrations of neurotransmitters including glutamate, glutamine, and GABA using LC-MS/MS in *Cln2^R207X^* cortices at 3 months which is immediately before the onset of spontaneous seizures. There were no significant changes in the concentrations of those neurotransmitters compared with wild type (Figure 8D).

**Figure 8.**
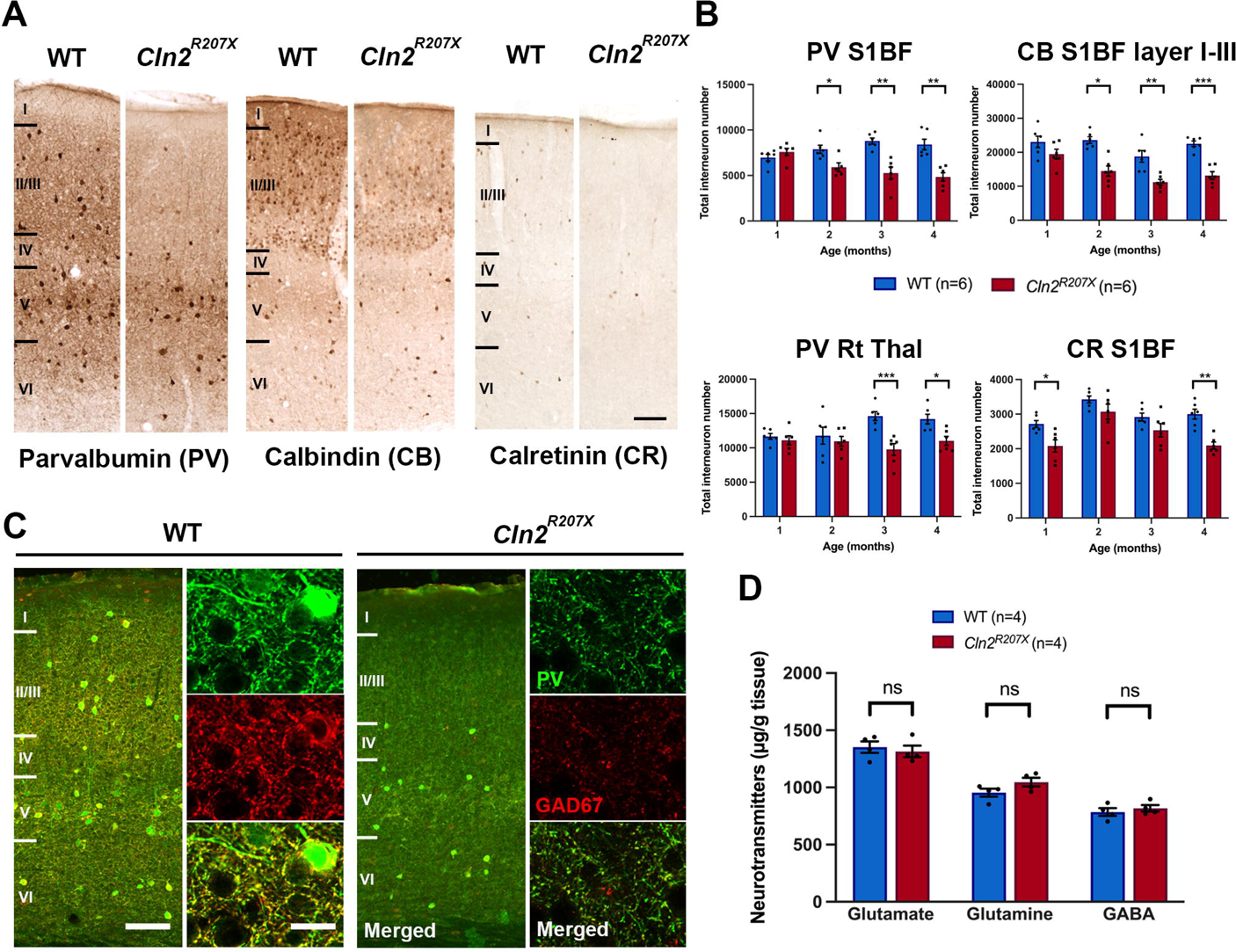
*Cln2^R207X^* mice show early GABAergic interneuron loss. (*A*) Immunoperoxidase staining for parvalbumin (PV), calbindin (CB), and calretinin (CR) show fewer of interneurons positive for these markers in the primary somatosensory cortex (S1BF) of *Cln2^R207X^* mice compared with WT mice at 4 months of age. Scale bar = 200 μm. (*B*) Unbiased stereological counts confirms the significant loss of PV- and CB-positive interneurons within S1BF in *Cln2^R207X^* mice (red bars) compared with WT mice (blue bars) as early as 2 months of age. Significant loss of PV-positive interneurons within reticular nucleus of thalamus (Rt Thal) at 3 months and loss of CR-positive interneurons in the S1BF at 1 and 4 months were also evident in *Cln2^R207X^* mice. Dots represent values from individual animals. \**P* < 0.05, \*\**P* < 0.01, \*\*\**P* < 0.001, \*\*\*\**P* < 0.0001, multiple t-test with Holm-Šídák correction. Values are shown as mean ± SEM (n = 6 mice per group). (*C*) Decreased immunoreactivity for PV (green) and GAD67 (red) in neuron fibers, especially within layers II-III in the S1BF in *Cln2^R207X^* mice at 3 months compared with age-matched WT mice detected by co-immunofluorescence staining. Images on the right side of each column are taken via confocal microscopy from within layers II/III. Scale bars = 20 μm (right) and 200 μm (left). (*D*) Measurement of neurotransmitters including glutamate, glutamine, and GABA in WT and *Cln2^R207X^* mouse cortex homogenates at 3 months by LC-MS/MS shows no significant difference between genotypes. One-way ANOVA with Bonferroni correction. Values are shown as mean ± SEM (n = 4 mice per group).

## Discussion

Previous CLN2 disease research has primarily focused on the development of therapies, which led to the approval of ERT with cerliponase alfa in the US and Europe in 2017 [8]. Although ERT slows CLN2 disease progression[9], cerliponase alfa is not curative and requires bi-weekly life-long administration. Alternative therapeutic approaches for CLN2 disease including viral gene therapy have been under preclinical development [42–44], but have not so far demonstrated efficacy upon clinical testing [45, 46]. In order to improve the therapeutic efficacy it is crucial to close the gap in knowledge of CLN2 disease pathophysiology at neuroanatomical and cellular levels. However, compared to the considerable efforts put in developing a therapy for this disorder, the fundamental mechanism(s) by which TPP1 deficiency causes such devastating neurodegeneration remains poorly investigated. This current multi-disciplinary study has not only revealed a distinct staging of neuropathology in forebrains and spinal cords of *Cln2^R207X^* mice, but also systematically defined their spontaneous seizure phenotype. This is a key presenting symptom in CLN2 patients, but until now has not been investigated in any animal model of CLN2 disease. Our findings also provide new insights to understand CLN2 disease pathophysiology and how it differs from other forms of NCL which may be used to improve the targeting of therapeutic strategies.

Early localized activation of both microglia and astrocytes that proceeds subsequent neuron loss is a neuropathological hallmark across multiple forms of NCL [10, 11]. Progressive microglial activation and astrogliosis are also present in *Cln2^R207X^* mice, and are most pronounced in the sensory corticothalamic system compared with other CNS regions. However, unlike other forms of NCL, astrogliosis in *Cln2^R207X^* mice did not coincide with the early onset of microglial activation at 1-2 months depending on the region. Instead, astrogliosis was relatively delayed and paralleled or immediately preceded the onset of Nissl-stained neuron loss at least one month later. Another marked difference from other forms of NCL is the regional specificity of neuropathology in *Cln2^R207X^* mice. Our data for both Nissl-stained and interneuron loss reveal the first detectable neuron loss to be in the cortex before subsequently occurring in the thalamus of *Cln2^R207X^* mice. This is in the opposite order to which these brain regions are affected in other forms of NCL where loss of thalamic relay neurons precedes neuron loss in the corresponding cortical target region [12–17]. Similarly, in contrast to the early involvement of the spinal cord compared to the forebrain in CLN1 [18, 19], we found that spinal pathology in *Cln2^R207X^* mice only starts after brain pathology is present. Moreover, unlike CLN1 disease [24], there was no significant loss of either Nissl-stained or interneuron loss in the hippocampus of *Cln2^R207X^* mice. Taken together our results reveal a novel regional pattern of neuropathological progression in CLN2 disease that is in stark contrast to other forms of NCL. This is especially the case compared to CLN1 disease, where the spinal cord, thalamus and cortex are affected in the opposite sequence [18, 19].

CD68 and GFAP are commonly used markers for microglial activation and astrogliosis, respectively, across many neurodegenerative diseases including the NCLs. However, there is emerging evidence suggesting a more complex and heterogenous signature of activated microglia and astrocytes, introducing the concept of M1/M2 microglia or A1/A2 astrocyte activation states along with their unique molecular markers [34–37]. Here, we found a delay of several months between the early onset of CD68-positive microglial activation and the late onset of elevated staining for MHC-II (M1 marker) and Clec7a (M2 or DAM) in *Cln2^R207X^* brains, suggesting progressive differences in the nature of microglial activation during CLN2 pathogenesis. In addition, our RT-qPCR evidence for changes in the expression of only a subset of A1/A2-specific astrocytic markers are in stark contrast to the pronounced typical A1-specific molecular signature transformation that occurs in *Cln1^-/-^* mice [47]. While the contribution of the heterogeneous subtypes of reactive astrocytes to neurodegeneration remains unclear, such data highlight the distinct molecular and functional nature of astrogliosis in CLN2 disease compared with CLN1 disease.

Our data revealed progressive changes in the expression of a restricted subset of cytokines and chemokines in the forebrain of *Cln2^R207X^* mice. The early elevation of IL- 33, IP-10 (CXCL10), and MIP-1α (CCL3) is shared in both *Cln2^R207X^* forebrains and *Cln1^-/-^* spinal cords [19], suggesting their potential involvement in the initial pathogenesis of both diseases, albeit in different regions of the CNS. IL-33 is an immediate indicator of tissue stress acting as an “alarmin,” predominantly expressed in astrocytes and oligodendrocytes in the CNS [48, 49]. Opposing roles for IL-33 have been suggested in different conditions with proposed pro-inflammatory roles in pain disorders and potentially anti-inflammatory roles in autoimmune encephalomyelitis, Alzheimer disease, and ischemic and traumatic CNS injury [50]. While the pathological role of IL-33 in *Cln2^R207X^* mice remains to be elucidated, our data suggest that early elevation of IL-33 may trigger downstream microglial-mediated neuroimmune responses, supported by evidence of predominant expression of both IP-10 (CXCL10) and MIP-1α (CCL3) in activated microglia [34,51–53]. Meanwhile, there were no changes in the expression of multiple cytokines and chemokines including IFN-γ and IL-10 that are significantly altered in *Cln1^-/-^* spinal cords [19]. A lack of change in the expression of these central pro- and anti-inflammatory molecules implies that CLN2 disease may be a less ‘neuroinflammatory’ disease than CLN1 disease.

Culturing both microglia and astrocytes derived from *Cln2^R207X^* mice enables us to compare morphological and functional alteration of those cell types with those from *Cln1^-/-^* and *Cln3^-/-^* mice. Unlike microglia derived from *Cln1^-/-^* and *Cln3^-/-^* mice [28, 29], *Cln2^R207X^* microglia appear morphologically normal under basal conditions and retain their capability to transform morphologically under pharmacological stimulation. In contrast, similar to astrocytes derived from *Cln3^-/-^* mice [28], *Cln2^R207X^* astrocytes appear morphologically indistinguishable from WT astrocytes under basal conditions, but do not elevate GFAP expression following pharmacological stimulation. Such *in vitro* results may reflect our histological data for a delayed onset of astrogliosis in *Cln2^R207X^* mice. Nevertheless, it will be informative to investigate whether *Cln2^R207X^* microglia and astrocytes exert a direct influence upon astrocytes and neurons using the similar co-culture techniques used in our previous studies [28, 29], or in microglial- and astrocytic-specific *Cln2^R207X^* mice.

Unexpectedly, gait abnormalities in *Cln2^R207X^* mice appeared only towards disease end stage, and to a remarkably lesser extent than those observed in *Cln1^-/-^* mice [19]. This may be related to the less severe and delayed onset of spinal cord pathology in *Cln2^R207X^* vs. *Cln1^-/-^* mice. These data are also unexpected given that severe gait decline is seen in patients with CLN2 disease [54]. This raises the possibility that *Cln2^R207X^* mice may die due to other causes, such as seizures, before their gait abnormality becomes severe enough to prevent them from accessing their food and water. The interictal EEG background abnormalities and pronounced spontaneous seizure phenotypes in *Cln2^R207X^* mice revealed by our longitudinal EEG recording are progressive in nature, and comparable with those in *Cln1^-/-^* mice and other genetic epilepsy models such as tuberous sclerosis complex 1 mice [13, 31]. However, a majority of *Cln2^R207X^* mice died shortly after seizures, and it will be important to determine whether these events may be related. Our EEG findings provide a novel, clinically relevant behavioral phenotype in *Cln2^R207X^* mice which will help us better understand the etiology of CLN2 disease, and also serve as a robust readout that can be used in future pre-clinical therapeutic studies.

Defining seizure phenotypes in *Cln2^R207X^* mice guided us to extend our pathological characterization of these mice to explore potential GABAergic interneuron deficits, which are implicated in contributing to both neurodegenerative diseases and epilepsy [40,41,55]. Our data reveal cortical interneurons are severely affected in *Cln2^R207X^* mice. Interestingly, no hippocampal interneuron loss was apparent, even at disease end stage. Within the cortex interneuron populations expressing different calcium binding proteins were affected equally in *Cln2^R207X^* mice, and were consistently more severely impacted in layers II-III than in deeper cortical layers. However, cortical interneuron loss invariably preceded that of Nissl-stained neurons in the same cortical region, or the loss of parvalbumin positive interneurons in the thalamic reticular nucleus. Although our EEG monitoring cannot precisely locate the anatomical source of seizure activity, our histological data suggest a cortical rather than hippocampal origin for seizures in *Cln2^R207X^* mice. Nevertheless, it is surprising that there was no significant change in the overall concentration of cortical neurotransmitters including GABA. This may reflect compensatory mechanisms to maintain total cortical GABA levels, or the inability of our analysis to distinguish between intracellular and extracellular levels of cortical neurotransmitters. It will be important to further investigate the nature of interneuron dysfunction and how this may contribute to epileptogenesis in *Cln2^R207X^* mice.

In conclusion, our data reveal phenotypes in *Cln2^R207X^* mice that differ markedly in their nature and timing compared to other mouse models of NCL. This includes the relative timing and distribution of glial activation and neuron loss, the relative onset of brain vs. spinal and cortical vs. subcortical pathology, and relatively late onset of gait abnormalities. Our data also reveal robust seizure phenotypes, which may be related to the pronounced loss of cortical interneurons. These data emphasize that while disease presentation may be broadly similar across forms of NCL, our detailed characterization has revealed phenotypes in CLN2 disease mice that are unique in many fundamental aspects. Our findings also pave the way for assessing the cellular specificity of TPP1 deficiency, which appears to differ from other forms of NCL.

## Authors’ contribution

JDC, KT, MW, and MSS conceived and designed the study. KT carried out the pathology experiments, tissue culture experiments, RT-qPCR, and statistical analysis on results from all experiments; EME performed gait analysis, SHW performed a part of immunohistochemistry, and NRR performed EEG recording. As acknowledged below, chemokine and cytokine analyses were performed by the core facility in the Bursky Center for Human Immunology and Immunotherapy Programs, Washington University School of Medicine and Barnes-Jewish Hospital in St. Louis, and LC-MS/MS were performed by the Washington University Metabolomics Facility, Washington University School of Medicine in St. Louis. HRN participated in mouse sample preparation and contributed to interpretation of histological data. JTD contributed to interpretation of gait analysis data. KT and JDC wrote the manuscript with input from all the authors. All authors read and approved the final manuscript.

## Acknowledgements

We thank the Washington University Center for Cellular Imaging (WUCCI) supported by Washington University School of Medicine, the Children’s Discovery Institute of Washington University and St. Louis Children’s Hospital, and the Foundation for Barnes-Jewish Hospital in St. Louis, for providing a Zeiss Axio Scan Z1 Fluorescence Slide Scanner, the Molecular Microbiology Imaging Facility at Washington University School of Medicine in St. Louis for providing a Zeiss LSM880 Confocal Laser Scanning Microscope with *Airyscan,* Alvin J. Siteman Cancer Center at Washington University School of Medicine and Barnes-Jewish Hospital in St. Louis, for the use of the Bursky Center for Human Immunology and Immunotherapy Programs, which provided the Luminex multiplex service, and the Washington University Metabolomics Facility at Washington University School of Medicine in St. Louis for providing the LC-MS/MS service. We acknowledge Dr. Diane Bender, Washington University in St. Louis for assistance with cytokine profile analysis and Dr. Xuntian Jiang, Washington University in St. Louis for assistance with neurotransmitter analysis. We also acknowledge Dr. Alison Barnwell for constructive comments on the manuscript.

## Ethics approval and consent to publication

All animal procedures were performed in accordance with National Institutes of Health (NIH) guidelines under protocols 2018-0215 and 21-0292 approved by the Institutional Animal Care and Use Committee (IACUC) at Washington University School of Medicine in St. Louis, MO.

## Data availability statement

The data sets analyzed in this study are available from the corresponding author upon reasonable request.

## Figure Captions

**Figure S1. *Cln2^R207X^* mice show altered expression of multiple markers of microglial activation and astrogliosis.** (*A*) Immunostaining for the ‘M1’ microglial marker MHC class II (MHC-II, green) and the ‘M2’ or disease-associated microglial (DAM) marker Clec7a (red) reveals increased immunoreactivity for these markers within the ventral posterior nuclei of thalamus (VPL/VPM) of *Cln2^R207X^* mice at 4 months of age. Scale bar = 200 μm. (*B*) RT-qPCR analysis shows limited expression of A1/A2 astrocyte-specific genes in *Cln2^R207X^* mouse forebrains compared with WT control mice at 3 and 4 months of age.

**Figure S2. *Cln2^R207X^* mouse forebrains show distinct neuroinflammatory changes.** Observed concentrations (pg/ml) of multiple cytokines and chemokines measured in forebrain homogenates reveal no significant change of IL-1β, TNF-α, IFN-γ, IL-6, IL-1α, IL-2, CXCL1, CRP, IL-10, IFN-α, IFN-β, and Eotaxin in *Cln2^R207X^* forebrains (red bars) compared with age-matched WT controls (blue bars). Dots represent values from individual animals. \**P* < 0.05, \*\**P* < 0.01, \*\*\**P* < 0.001, \*\*\*\**P* < 0.0001, multiple t-test with Holm-Šídák correction. Values are shown as mean ± SEM (n = 6 mice per group).

**Figure S3. Primary microglia and astrocytes derived from *Cln2^R207X^* mice show abnormal morphology and response to pharmacological stimulation.** (*A*) Primary microglia derived from WT and *Cln2^R207X^* neonatal cortices are visualized with Iba-1 (green) for morphological analysis with and without treatment with LPS. Scale bar = 50 μm. (*B*) Under basal condition, there is no difference in aspect ratio and roundness between *Cln2^R207X^* and WT microglia. Following LPS treatment, aspect ratio and roundness increase in both *Cln2^R207X^* and WT microglia. The box extends from the 25^th^ to 75^th^ percentiles with the line in the middle of the box showing the median, and the whisker extend from the smallest and the largest values (n > 100 cells per group). (*C*) Primary astrocytes derived from WT and *Cln2^R207X^* neonatal cortices are immunostained for astrogliosis marker glial fibrillary associated protein (GFAP, red) with α-tubulin (green), both with and without LPS/IFN-γ stimulation. Scale bar = 50 μm. (*D*) The percentage of GFAP-positive astrocytes was increased by LPS/IFN-γ stimulation for WT astrocytes, but not for *Cln2^R207X^* astrocytes. Values are shown as mean ± SEM (n = 3 mice per group). \**P* < 0.05, \*\**P* < 0.01, \*\*\**P* < 0.001, \*\*\*\**P* < 0.0001, one-way ANOVA with Bonferroni correction.

**Figure S4. *Cln2^R207X^* mice show no neuron loss in the hippocampus.** Unbiased stereological counting reveals no change in the number of Nissl-stained neurons within CA1, PV-positive interneurons within the stratium oriens, CA1, CA2/3, and dentate gyrus, CB-positive interneurons within stratum oriens and radiatum, and CR-positive interneurons within CA1/2/3, radiatum, and hilus of *Cln2^R207X^* mouse hippocampi compared with age-matched WT control. Dots represent values from individual animals. \**P* < 0.05, \*\**P* < 0.01, \*\*\**P* < 0.001, \*\*\*\**P* < 0.0001, multiple t-test with Holm-Šídák correction. Values are shown as mean ± SEM (n = 6 mice per group).

**Figure S5. Neonatal *Cln2^R207X^* mice show storage material accumulation in the brain.** Immunostaining for subunit c of mitochondrial ATP synthase (SCMAS, green) reveal a widespread and pronounced SCMAS accumulation in *Cln2^R207X^* mouse forebrains at postnatal day 1 (P1). S1BF; primary somatosensory cortex, VPL/VPM; ventral posterior nuclei of thalamus, DLG; dorsolateral geniculate nucleus. Scale bar = 200 μm.

