## Supplementary figures and images for "Cortical interneuron loss and seizure generation as novel clinically relevant disease phenotypes in *Cln2^R207X^* mice"

### Supplemental Fig 1

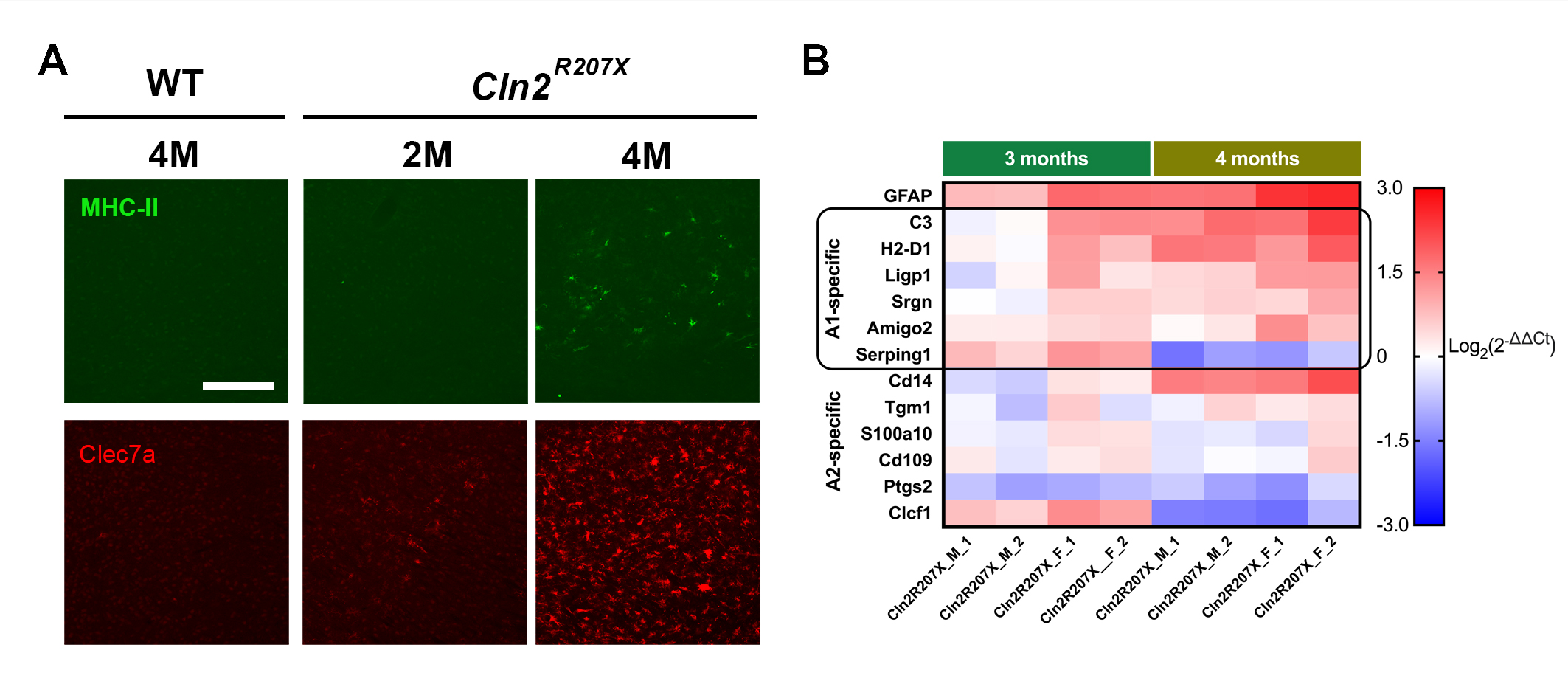

### Supplemental Fig 2

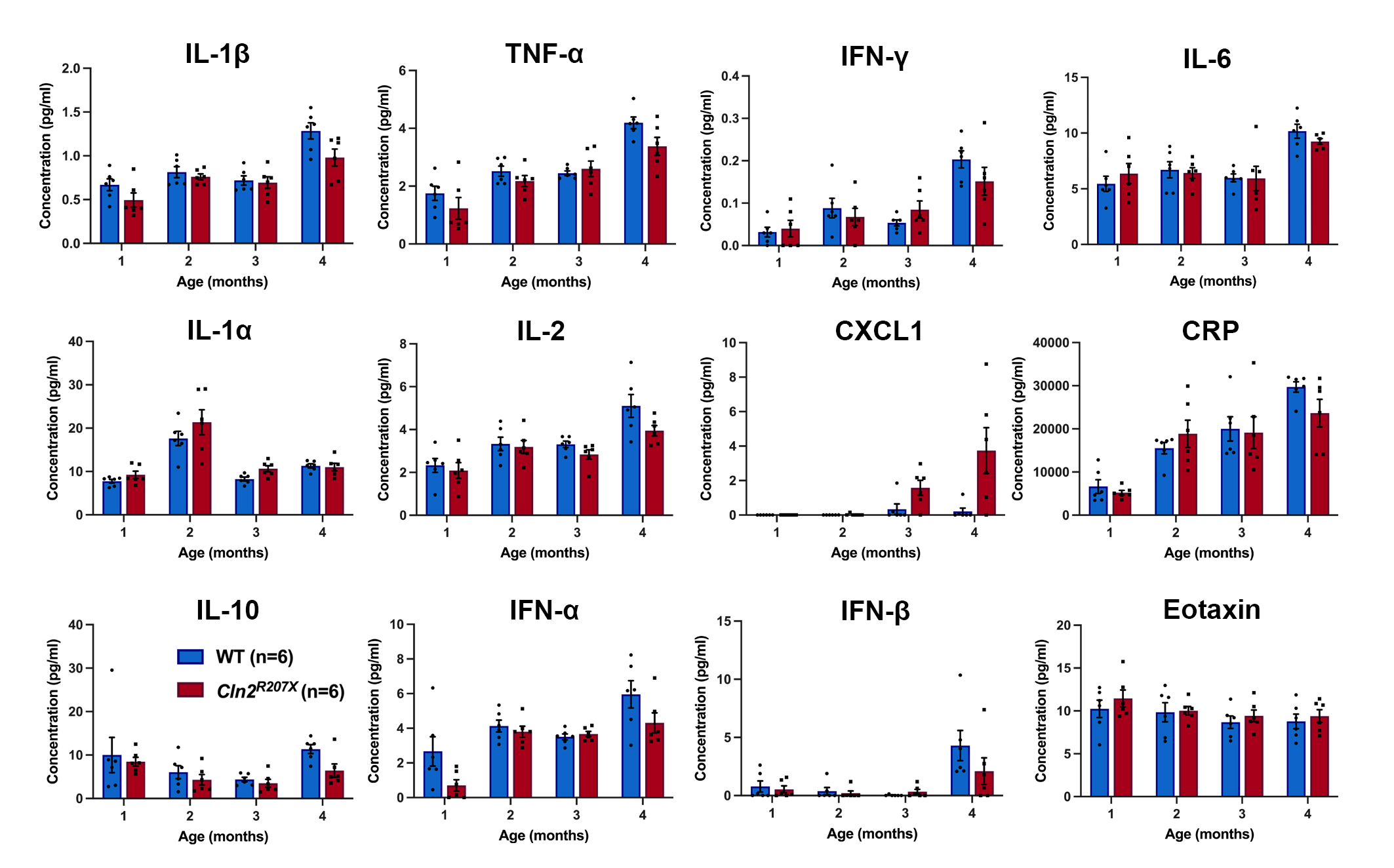

### Supplemental Fig 3

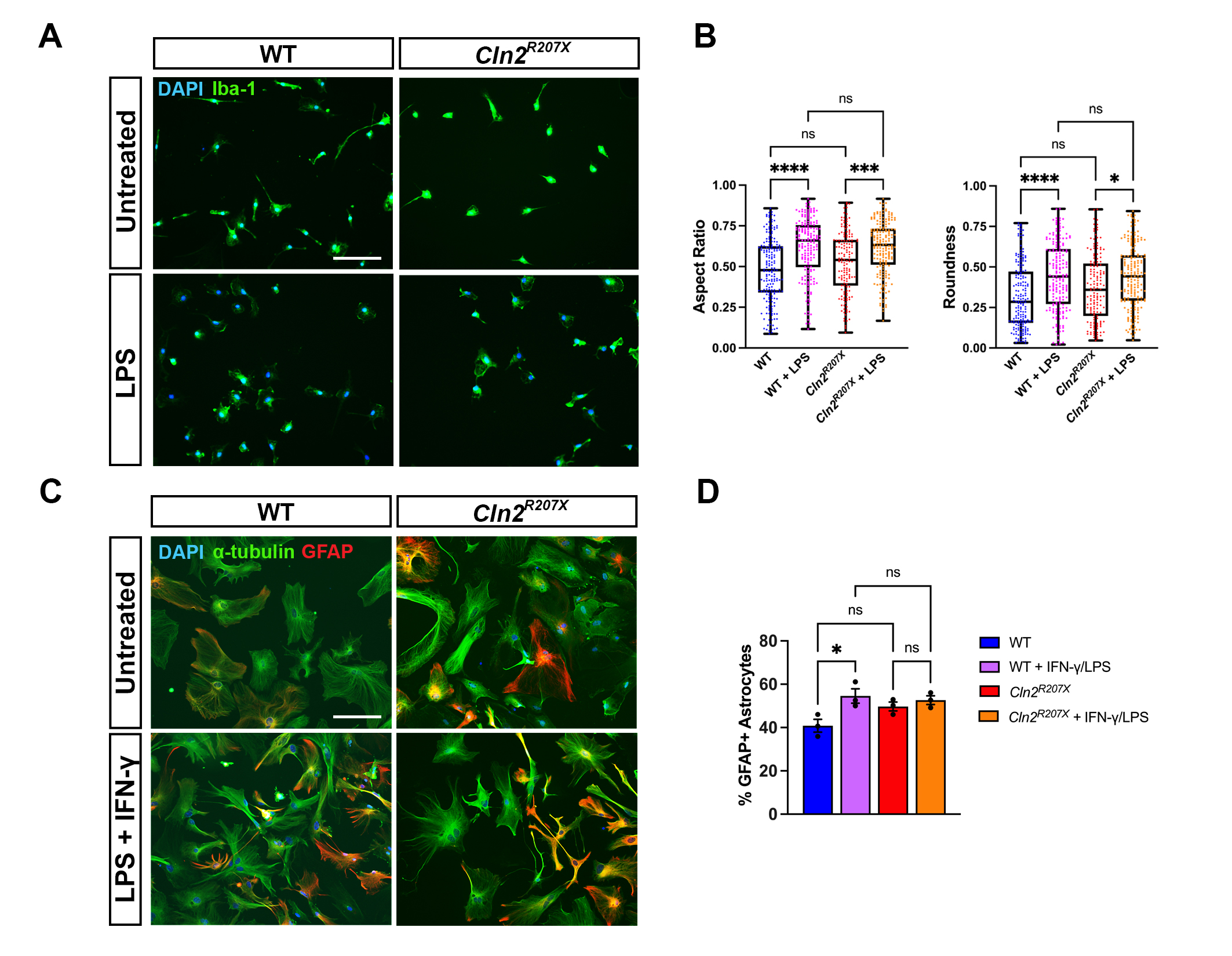

### Supplemental Fig 4

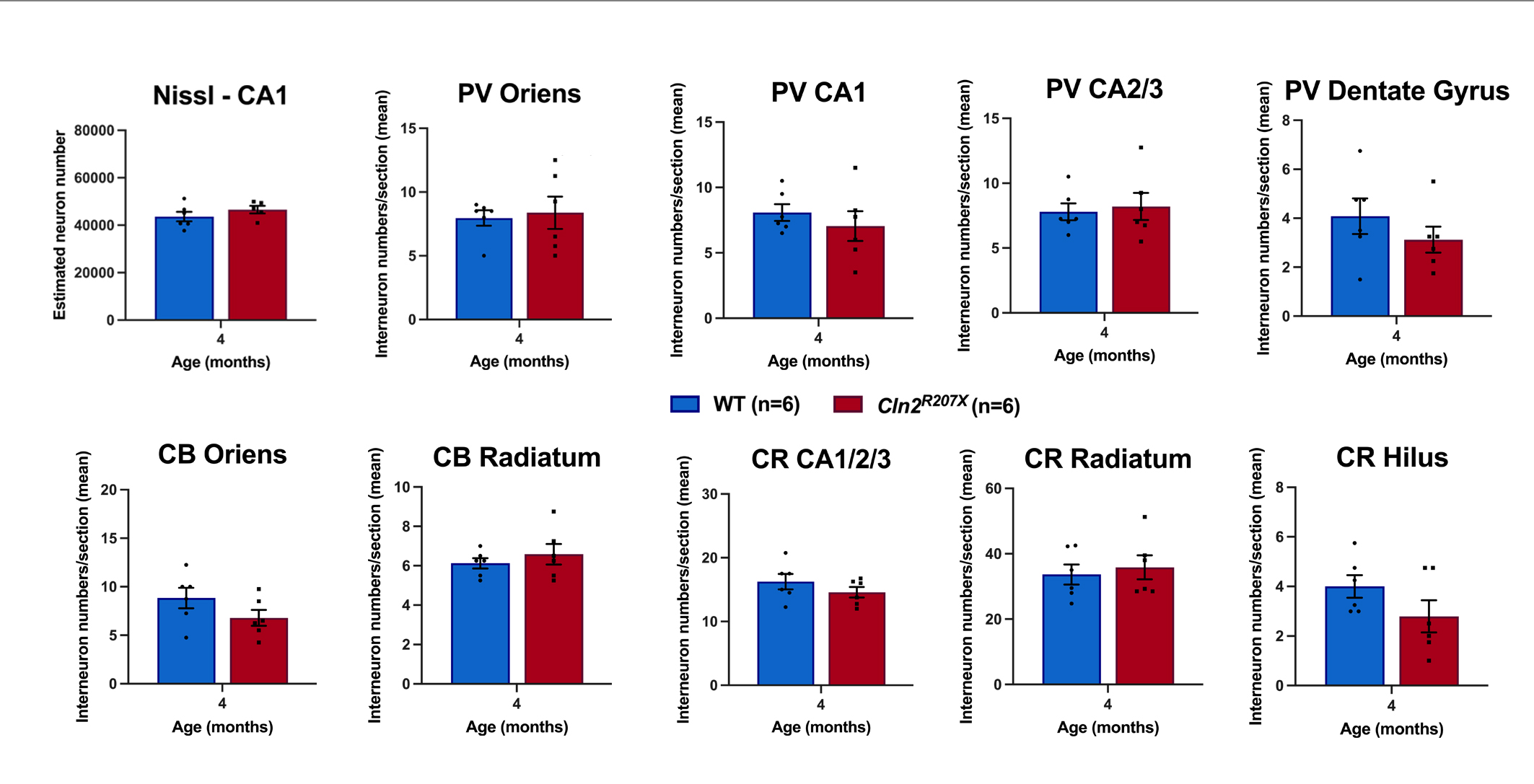

### Supplemental Fig 5

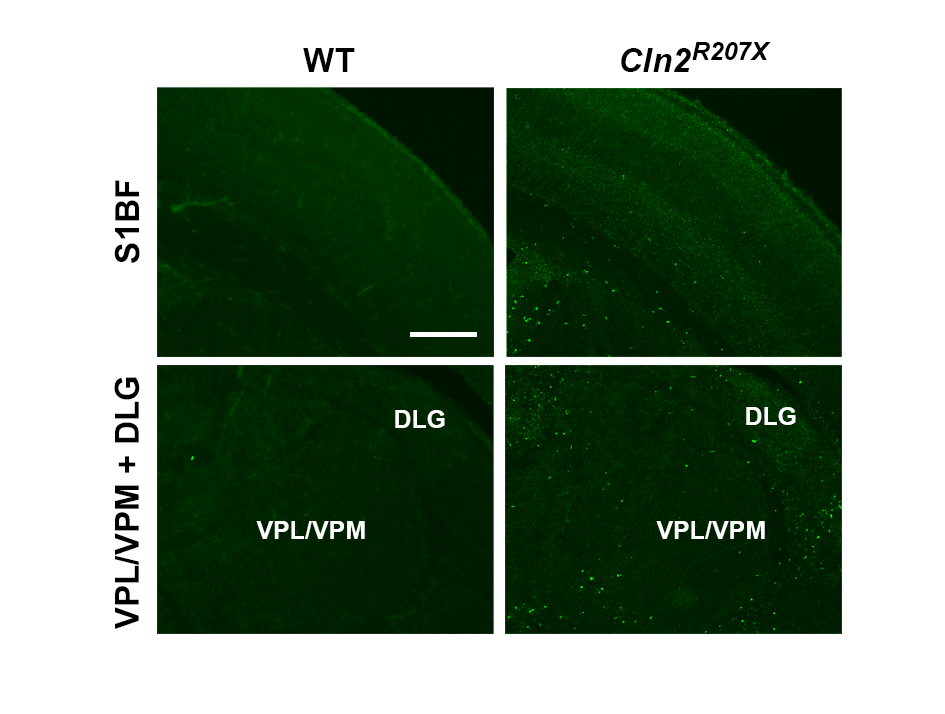
